# Does male mate choice select for female colouration in a promiscuous primate species?

**DOI:** 10.1101/2020.12.08.415810

**Authors:** Lucie Rigaill, Cécile Garcia

## Abstract

The traditional view of sex roles and sexual selection emphasises the evolution of male ornaments as a result of female mate choice and male-male competition. Female ornaments are now receiving more attention, although their roles in mating decision are still less well understood, especially considering cases in which colourful ornaments are expressed by both sexes. In this study, we analysed whether female skin colouration (luminance and redness of the face and hindquarters) influenced male mate choice and sexual behaviours in relation to intra-cycle (cycle phase), inter-cycle (number of consecutive cycles, conceptive nature of the cycle), and inter-individual (age, social rank, weight, and parity) variation in a captive population of Japanese macaques (*Macaca fuscata*). Males did not preferentially choose darker/redder females. Moreover, males did not appear to use female skin colouration to apportion their mating efforts on the most fertile period of the menstrual cycle or during cycles that lead to conception, or to discriminate between females. To our knowledge, our study is among the few to report a lack of male choice for female colouration in a species where both sexes potentially display ornamentation. While female colouration appeared to contain information about intra-cycle, inter-cycle, and inter-individual variation in fecundity, this study further demonstrates that this trait may not have been sexually selected and that males mated regardless of such variation across females. This study adds to the growing research on the role and evolution of female colouration in the context of sexual signalling and mate attraction.

## Introduction

Mate choice is a crucial process in reproduction. Individuals have to simultaneously choose and avoid potential mates based on the expected benefits (e.g., good genes, social support, reproductive success) and costs (e.g., disease transmission, energy costs, intra-sexual competition) associated with mating (Beltran-Bech & Richard, 2014; Møller et al., 1999; Thrall et al., 2000; Trivers, 1972). The traditional view of sex roles and sexual selection has emphasised the importance of female mate choice and male competition, and, as a consequence, the function of male ornaments has been well studied across taxa (Andersson & Simmons, 2006, 2006; Cunningham & Birkhead, 1998; Fitzpatrick & Servedio, 2017). Perhaps partly as a result, the role of male mate choice on the evolution of female traits has been relatively less investigated. However, males can also face costs associated with mating in the form of energetically demanding courtship displays, mate guarding, and the production of ejaculates (Balshine et al., 2002; Dewsbury, 1982; Harshman & Zera, 2007; Lukas & Clutton-Brock, 2014). Similarly, female competition can also arise when females vary in reproductive quality or when the number of females exceeds male mating or mate-guarding capacities such as in cases with a high ovarian cycle synchrony or with a female-biased operational sex ratio, i.e., the ratio of the number of fertile adult males to the number of potentially fertile females in a group at a given time (Amundsen, 2000; Bonduriansky, 2001; Edward & Chapman, 2011; Kvarnemo & Ahnesjo, 1996). Thus, male mate choice or mutual mate choice can evolve in some species to balance the ratio of costs/benefits inherent in mating (Clutton-Brock, 2007; Fitzpatrick & Servedio, 2018; Johnstone et al., 1996; Kokko et al., 2003).

There is increasing evidence that males discriminate among possible mates in species with traditional and reversed-sex roles and with different mating systems (from monogamy to polygynandry). Males may choose older females that have already successfully reproduced and raised offspring (spiny-footed lizards, *Acanthodactylus erythrurus*: Belliure et al., 2018; chimpanzees, *Pan troglodytes*: Muller et al., 2006; brown widow spiders, *Latrodectus geometricus*: Waner et al., 2018). Males may also choose females of higher social ranks compared with lower-ranking ones. This may be particularly adaptative in species where social rank is maternally inherited: higher-ranking offspring may achieve better condition and higher survival than lower-ranking offspring (e.g., mandrills, *Mandrillus sphinx*: Setchell & Wickings, 2006). Males may also choose to mate with unmated vs. mated females to reduce sperm competition and increase their reproductive success (house crickets, *Acheta domesticus*: Assis et al., 2017; wolf spiders, *Schizocosa ocreata*: Roberts & Uetz, 2005). To mate selectively, males may thus rely on female signals and/or express preferences for female traits that correlate with reproductive quality, such as pheromones (widow spiders, *Lactrodectus hesperus*: Baruffaldi & Andrade, 2015; humans, *Homo sapiens*: Gildersleeve et al., 2012; wolf spiders: Roberts & Uetz, 2005), body size (long-tailed dance flies, *Rhamphomyia longicauda*: Funk & Tallamy, 2000; LeBas et al., 2003; stalk-eyed flies, *Diasemopsis meigenii*: Harley et al., 2013; rainbow darters, *Etheostoma caeruleum*: Soudry et al., 2020; hermit crabs, *Pagurus middendorffii*: Wada et al., 2011), or vocal pitch (humans: Pisanski et al., 2018).

Among female traits, colourful ornaments have received more attention than others. In some species, females express sex-specific colourful ornaments whose colour and/or size are attractive to males. Such colourful ornaments are likely to be sexually selected traits as they correlate with indices of fecundity, such as sexual or egg maturity, readiness to spawn, brood size, or offspring condition (two-spotted gobies, *Gobiusculus flavescens*: Amundsen & Forsgren, 2001; collared lizards, *Crotaphytus collaris*: Baird, 2004; cichlid fishes, *Pelvicachromis taeniatus*: Baldauf et al., 2011; blue crabs, *Calinnectes sapidus*: Baldwin & Johnsen, 2012; spiny-footed lizards: Belliure et al., 2018; cichlid fishes, *Mikrogeophagus ramizeri*: LaPlante & Delaney, 2020; pygmy halbeaks, *Dermogenys collettei*: Ogden et al., 2020; striped plateau lizards, *Sceloporus virgatus*: Weiss, 2002, 2016). Comparatively, studies on the role of non-sex-specific colourful ornaments (i.e., ornaments expressed by both sexes) have yielded a contrasting picture of the extent to which these ornaments modulate mate choice. Female colouration may indeed represent a genetically correlated response of selection for male traits with no clear adaptive value in females. Males may thus express no preference for such traits (three spined sticklebacks, *Gasterosteus aculeatus*: Wright et al., 2015) or preference may be merely explained by sensory exploitation rather than sexual selection (sockeye salmons, *Oncorhynchus nerka*: Foote et al., 2004). However, there is some evidence that males can show preferences toward non-sex-specific colourful ornaments that reflect female reproductive state (rhesus macaque, *Macaca mulatta*: Dubuc et al., 2009; Higham et al., 2010, 2011; agamid lizards, *Ctenophorus ornatus*: LeBas & Marshall, 2000) or reproductive success (blue tits, *Cyanistes caeruleus*: Doutrelant et al., 2008, 2012; Mahr et al., 2012; rhesus macaques: Dubuc, Winters, et al., 2014; rock sparrows, *Petroni petronia*: Pilastro et al., 2003; Griggio et al., 2009, 2005; brown boobies, *Sula leucogaster*: Montoya et al., 2018; blue-footed boobies, *Sula nebouxii*: Torres & Velando, 2005). Therefore, in these species, female colourful ornaments can have a crucial impact on the mating strategies and reproductive success of both males and females.

Here, we investigate the potential role of female colouration on male mate choice in a promiscuous primate species, the Japanese macaque (*Macaca fuscata*). Female mate choice has been suggested as an important process modulating mating strategies in this species (Fujita, 2010). However, this does not mean that males should not choose or discriminate among potential mates. While their parental investment is mostly restricted to gamete contribution, male Japanese macaques deal with high costs associated with mating. Japanese macaques are seasonal breeders and multi-mount ejaculators and thus invest a non-negligible portion of their daily energy expenditure for somatic maintenance and mating itself (Fooden & Aimi, 2005; Matsubara, 2003; Thomsen et al., 2006). Like males of other species, male Japanese macaques also face physical and physiological costs associated with extended consortships and mate-guarding, reduced feeding activity, and male-male aggression leading to increased vigilance, injuries, or chronic stress (e.g., in several mammal species including primates: Alberts et al., 1996; Girard-Buttoz et al., 2014; Higham & Maestriperi, 2014; Lukas & Clutton-Brock, 2014; Thompson & Georgiev, 2014). Males would thus benefit from discriminating among females to focus their mating effort on mates that are fertile and of “good quality”. In Japanese macaques, both sexes express red skin colouration (luminance: how light/dark the skin appears, redness: at least partially reflecting the colour of the blood itself) of the face and hindquarters that peaks during the mating season with sex hormones (Fooden & Aimi, 2005). Recent studies provided more insight into the potential signalling function of female colouration in this species and characteristics that males may be selecting for. Hindquarter colouration may signal the intra-cycle probability of ovulation while concealing its exact timing and face colouration may indicate early pregnancy (Rigaill et al., 2015, 2019). Hindquarter colouration may also contain information about the inter-cycle probability of conception (Rigaill et al., 2019). Lastly, there is some evidence that skin colouration may reflect inter-individual variation (e.g., social rank, weight, but not parity, Rigaill et al., 2017, 2019). Pflüger and colleagues (2014) showed that males increased attention toward an experimentally intense red-coloured version of an unknown female face. Thus, female colouration may be a sexually selected trait involved in mate choice in Japanese macaques if males appropriately use this colourful trait to inform their mating decisions. Yet, to our knowledge, no study has tested this hypothesis in a non-forced-choice discrimination situation.

Our study thus aims at identifying the potential role of female Japanese macaque face and hindquarter colouration (luminance and redness) on the likelihood that a male will engage or maintain sexual interactions with a female. If female colouration acts as a fertility signal modulating mate choice, we hypothesised that males should display more sexual behaviours toward darker/redder (i.e., stronger signal) females. We further hypothesised that the relationship between male response and female colouration should vary according to a) intra-cycle (timing of the fertile phase), b) inter-cycle (number of consecutive cycles and probability of conception across cycles), and/or c) inter-individual (age, social rank, weight, and parity) variation. If so, we predicted a positive relationship between male sexual behaviours and darker/redder female colouration a) when the likelihood of ovulation is the highest (i.e., during pre- and fertile phases vs. the post-fertile phase), b) during cycles that lead to conception (i.e., during conceptive cycles and the first cycles of a given mating season), and/or c) with older, higher ranking, heavier, and possibly multiparous females.

## Methods

### Subjects and housing

We collected data during the 2011-2012 mating season, from early November to late January, from a captive population of Japanese macaques living in a 1,210 m^2^outdoor enclosure at Kyoto University Primate Research Institute (KUPRI, Inuyama, Japan). The group was composed of 39 individuals consisting of 13 adult females (> five years, mean ± SD = 10.64 ± 6.84 years, range = 5-27), 6 sexually immature females (2.79 ± 0.81 years, range = 1-4), 3 adult males (> six years, 10.67 ± 3.06 years, range = 8-14), 12 sexually immature males (2.99 ± 1.05 years, range = 1-4), and 5 infants less than one year old. All adult and sexually active females and males in this group were included in this study. The ratio of adult males to adult females is similar to other captive and wild populations from studies of sexual behaviours and mating strategies (Fooden & Aimi, 2005; Fujita, 2010). We did not include subadult males/females to avoid confounding factors due to maturational age. Females were naturally cycling, i.e., with no hormonal contraceptive treatment. Female age, weight at the beginning of the mating season (to the nearest 100g), and parity (total number of infants born divided by the number of years post-adulthood) were provided by the Center for Human Modeling Research of KUPRI. Animals were fed twice daily between 11:00 AM and 12:00 PM and between 4:00 PM and 5:00 PM. Water was supplied *ad libitum*. The Center for Human Evolution Modeling Research of the Primate Research Institute reviewed and approved our research protocol in agreement with the Guidelines for the Care and Use of Nonhuman Primates of the Kyoto University Primate Research Institute.

### Behavioral observations

One observer collected behavioural data using focal animal sampling distributed from 7:00 AM to 5:00 PM, 7 days a week for 3 months. We used a female-centered approach because of the need to assess female trait variation in order to address the question of male responses. We followed each focal female 2 hours per week in 4 blocks of 30-min continuous focal samples equally distributed across days and females. During focal observations, we recorded all social and sexual behaviours between males and females, along with the direction of the behaviour between the focal female and any identified adult male. We recorded all occurrences of female presentations, female approaches, female mounts, female vocalisations (oestrus and copulation calls), male approaches, male holding behaviours, male inspections of female genitals (i.e., visual and/or tactile and/or olfactory inspections of the female genital area), and male mounts (i.e., ejaculatory and non-ejaculatory). The number of observed records for each behaviour is given in Table 1. Female and male social ranks were assessed by transcribing agonistic interactions for which a clear win/loss outcome was identified into an agonistic interaction matrix. We then calculated the Normalized David’s Score (NDS) to assess female social rank positions (de Vries et al., 2006). Given that there were only three adult males in the group, male ranks were assessed as alpha (winning all agonistic interactions), beta (defeating all males except the alpha male), and gamma (losing all agonistic interactions) directly from the agonistic interaction matrix.

**Table 1.** Total number of female behaviours, male behaviours, and female vocalisations recorded.

|  | Sum |
| --- | --- |
| *Female behaviours* |  |
| Presentations | 10 |
| Approaches | 86 |
| Mounts | 86 |
| *Male behaviours* |  |
| Approaches | 153 |
| Holding behaviors | 54 |
| Genital inspections | 39 |
| Mounts | 75 |
| *Female vocalisations* | 30 |

### Female colouration

One experimenter took digital images of female face and hindquarters every two days in the morning between 11:00 AM and 12:00 PM. We used a Canon EOS 350D camera with an 8 megapixel CMOS censor and an EF28–135 mm f/3.5–5.6 IS USM lens. Images were standardized by daily manual setting of the white balance using an X-Rite White Balance Card and a Gretag X-Rite Color Checker (GretagMacbeth ColorChecker). For each female’s photo, we assessed colour from the whole face and hindquarter areas excluding the eyes, nose, forehead, and tail. A technician who was blind to females’ identity and characteristics extracted reflectance spectra using Colourworker software (Chrometics Ltd. Website, available: http://www.chrometrics.com). These spectra were converted into quantal catches, i.e., the stimulation of Japanese macaque photoreceptors (Rigaill et al., 2019) using equations and macaque cone ratio and spectral sensitivity values given elsewhere (Higham et al., 2010; Stevens et al., 2009). Based on the quantal catch data for both face and hindquarter colouration, we calculated luminance (how light/dark the skin appears) and redness (variation in chromatic parameters) data.

### Female reproductive status

To determine each female’s reproductive status, we collected a total of 381 faecal samples (mean ± SD per female = 29.31 ± 3.17, range = 24-34) on average every 1.59 days (range = 0-4 days) to accurately assess the ovarian cycle (Hodges & Heistermann, 2011). We collected faecal samples in their entirety immediately after voiding, with most samples collected between 8:00 AM and 12:00 PM. All samples were then stored at −20°C until processing. Faecal samples were analysed for pregnanediol-3-glucuronide (PdG) using enzyme immunoassays as previously described by Garcia et al. (2009) to assess the presumed day of ovulation and the periovulatory period (15-day period around the estimated ovulation) of each ovarian cycle. The onset of the luteal phase was determined as the sample with a faecal PdG concentration which was at least two standard deviations greater than the mean PdG concentration of the 3–4 preceding baseline values (Hodges & Heistermann, 2011). We determined the 2-day window for ovulation as days −2 and −3 relative to the faecal PdG rise. The fertile phase was defined as a period of five days (covering a two-day window for ovulation plus three preceding days to account for sperm longevity in the reproductive tract (Behboodi et al., 1991). The five-day period preceding the fertile phase represented the pre-fertile phase, while the five-day period following the fertile phase was defined as the post-fertile phase. We recorded a total of 24 ovarian cycles (mean ± SD per female = 2.0 ± 0.83, range = 1-3) and the majority of these cycles were non-conceptive (19/24). One female failed to cycle during the study and was thus excluded from the analysis.

## Data analysis

We restricted our analyses to the pre-fertile, fertile, and post-fertile phases. We used 173 focal observations with matching female photo (i.e., focal and photo on the same observation day) for the analyses (mean ± SD = 7.53 ± 1.21 per female and per cycle). We built our dataset based on the 30-min focal observations and attributed one line to each focal female – male dyad per focal observation (i.e., three lines per focal female), which was the unit of analysis.

We carried out our analyses in R version 3.3.3. Before running the statistical analyses, we checked for correlations between our variables of interest. Female weight correlated with parity (rs = −0.50, P < 0.001) and age (rs = 0.73, P < 0.001). Parity correlated with age (rs = −0.42, P < 0.001). Parity also weakly correlated with female social rank (rs = −0.25, P < 0.001). We thus excluded female age and weight from our analyses. None of the female behaviours (i.e., presentations, approaches, and mounts) were correlated with each other (all P > 0.05). None of the male behaviours (i.e., approaches, holding behaviours, genital inspections, and mounts) were correlated with each other (all P > 0.05).

We aimed at testing whether the likelihood of a male behavioural response (binary variable: 1 = occurrence, 0 = non-occurrence) toward a female was predicted by female colouration, i.e., whether males where more likely to engage in or maintain sexual interactions with darker/redder females. The little occurrences of copulatory event alone prevented us from analysing the effect of female colouration on this behaviour only. We thus computed a binary response variable that corresponded to a male behavioural response as the occurrence of at least one of the behaviours involved in establishing and maintaining consortship that were displayed by the interacting male of the dyad (i.e., approaches, holding behaviours, genital inspections, and copulations). Similarly, we also computed a binary variable that corresponded to a female behavioural display (hereafter, *female behaviours*), as the occurrence of at least one of the behaviours involved in establishing and maintaining consortship that were displayed by the interacting female of the dyad (i.e., presentations, approaches, and mounts).

We built a series of generalized linear mixed models with a binomial error structure and logit link function with the function glmer from the lme4 package (Bates et al., 2015) to investigate the relationship between male behavioural response (binary response variable) and female hindquarter and face luminance and redness (continuous predictor variables). Inspection of the cumulative distribution functions revealed good fits to the normal distribution for hindquarter luminance and to the lognormal distribution for face luminance, face redness, and hindquarter redness. We first constructed three different models to test the relationship between male behaviours and female face colouration (luminance and redness), male behaviours and female hindquarter colouration (luminance and redness), and male behaviours and female face and hindquarter colouration (luminance and redness). To account for a) intra-cycle, b) inter-cycle, and c) inter-individual effects on the potential relationship between female colouration and male response, our three models included interactions between female colouration and a) the cycle phase (categorical variable, three levels: pre-fertile, fertile, and post-fertile phases), b) the number of consecutive cycles (continuous variable) and the conceptive nature of the cycle (binary variable: 1 = conceptive, 0 = non-conceptive cycles, hereafter *conception*), and c) female social rank (continuous variable) and parity (continuous variable). We also included female behaviours (binary variable) and male social rank (categorical variable, three levels: alpha, beta, gamma) as covariates to account for the potential effects of female direct solicitation and male social attributes on mating decisions. Finally, the date of the focal, female identity, and male identity were included as random factors. We also constructed a null model in which the predictor variables and all covariates were removed but the random effects structure was maintained. All continuous variables were standardized to mean = 0 and SD = 1 prior to modelling to improve model performance and interpretability (Harrison et al., 2018). We checked for variance inflation (GVIF^(1/(2xDf))^< 3) using the car package (Fox & Weisberg, 2019); we excluded the number of consecutive cycles from the models due to variance inflation. We ensured that all relevant model assumptions (homogeneity of the residuals and stability of estimates) were met by visually inspecting histograms of the residuals and plots of residuals against fitted values.

We used an information-theory approach to objectively compare and rank the four candidate models (i.e., null model, face colouration model, hindquarter colouration model, and face and hindquarter colouration model) in terms of how well they fitted the existing data. This method allowed us to assess the likelihood that one or more models among the candidates is/are best supported by the data (Burnham et al., 2011; Symonds & Moussalli, 2011). We used the function model.sel of the MuMIn package (Bartoń, 2020) to rank models based on their respective Akaike’s information criterion corrected for small sample size (AICc values). We considered models with ΔAICc (the difference in AICc values between the smallest AICc and the other AICc values) below four as competitive models with the plausibly best model in the set of candidate models (Burnham & Anderson, 2002). We reported the weight of the models which indicates to what extent one candidate model is more likely than another to provide a reasonable explanation of the variance in the data, along with the evidence ratio (ER) which indicates the extent to which the higher-ranked model is more parsimonious than another candidate model. We also reported the marginal (fixed effects alone) and conditional (fixed effects and random structures) coefficients of determination (R^2^m and R^2^c) of all models using the function r.squaredGLMM (Nakagawa & Schielzeth, 2013). We extracted weighted parameter estimates (β), standard errors (SE), and 95% confidence intervals (95% CIs) of model intercepts and predictors from conditional averaging of the 4 candidate models (function model.avg).

## Results

Among the candidate models, the model including female hindquarter colouration (luminance and redness) and covariates had the highest weight (0.998) and more than 499 times more empirical support than the competing models (face colouration: ΔAICc = 12.3; face and hindquarter colouration: ΔAICc = 17.6; null model: ΔAICc = 49.8, Table 2). The fixed and random effects explained around 28% of the observed variance in the occurrence of male behavioural responses.

**Table 2.**
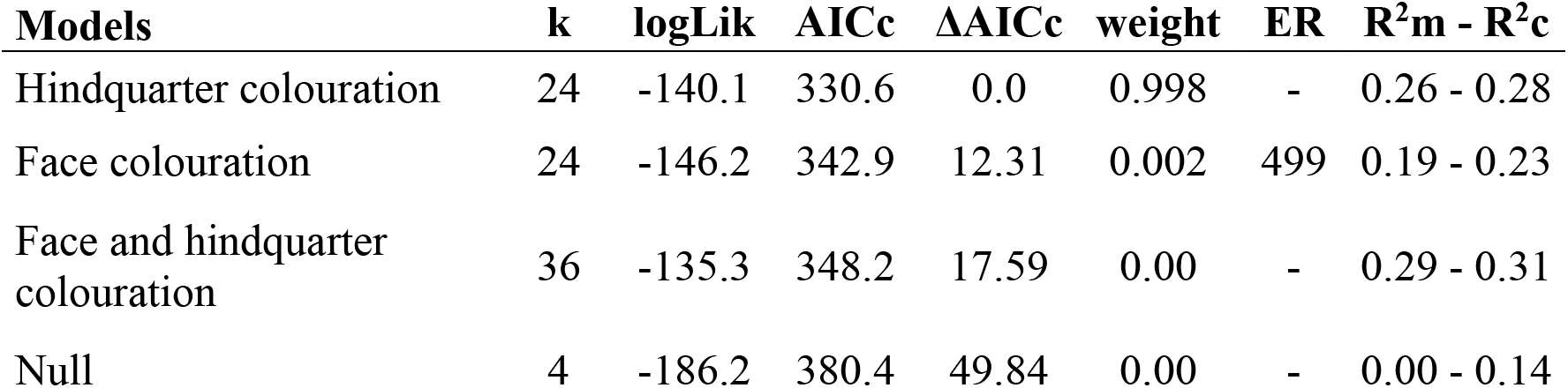
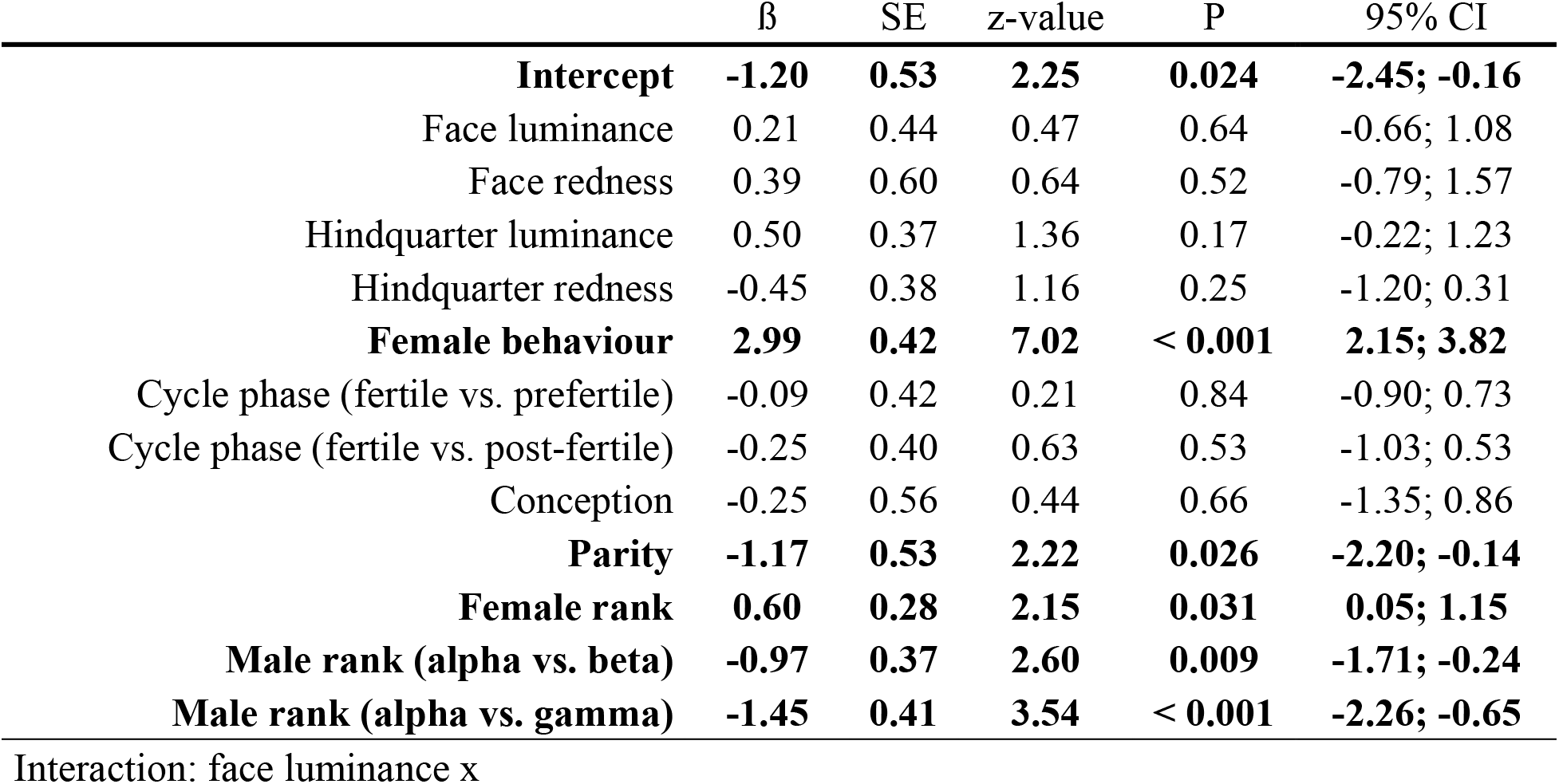
Characteristics of candidate models. *k*: number of predictors, *AICc*: corrected Akaike’s information criterion, *ΔAICc*: AICc difference between higher-ranked models and other candidate models., *weight*: model probabilities, *ER*: evidence ratios, *R^2^m – R^2^c*: marginal and conditional coefficients of determination. Candidate models are presented from higher-ranked to lower-ranked ones.

There was no effect of cycle phase (fertile vs. prefertile and post-fertile phases) on the interaction between male behavioural responses and female colouration (luminance and redness of the face and hindquarters, all P > 0.50, Table 3, Figure 1A). There was a slight tendency for a higher probability of male behavioural responses when hindquarter luminance was high in non-conceptive vs. conceptive cycles (ß = -0.96 ± 0.52 SE, P = 0.068, 95% CI =-1.98; 0.07, Table 3, Figure 1B). In other words, our results suggest that during conceptive cycles, there was a higher probability of male behavioural responses when the signal was strong (lower luminance = darker hindquarters). However, female face and hindquarter colouration *per se* did not explain variation in the occurrence of male behavioural responses (luminance and redness, all P > 0.05, Table 3). Similarly, the likelihood of observing male sexual behaviours was not influenced by cycle phase or the conceptive nature of the cycle (all P > 0.50, Table 3).

**Table 3.** Model averaged parameters estimates (ß) ± adjusted standard errors (SE), z-value, probability (P), and confidence intervals (95% CIs) from conditional averaging of all candidate models. For each categorical comparison, the first level indicate baseline. Results for which CI does not include zero are presented in bold, trends are presented in italic.

|  |  |  |  |  |  |
| --- | --- | --- | --- | --- | --- |
| Cycle phase (fertile vs. prefertile) | -0.43 | 0.47 | 0.91 | 0.36 | -1.34; 0.49 |
| Cycle phase (fertile vs. post-fertile) | -0.25 | 0.47 | 0.52 | 0.60 | -1.18; 0.68 |
| Conception | -0.50 | 0.47 | 1.07 | 0.28 | -1.42; 0.42 |
| Parity | 0.25 | 0.41 | 0.62 | 0.53 | -0.55; 1.06 |
| Female rank | 0.14 | 0.22 | 0.66 | 0.51 | -0.28; 0.57 |
| Interaction: face redness x |  |  |  |  |  |
| Cycle phase (fertile vs. prefertile) | 0.22 | 0.54 | 0.41 | 0.68 | -0.84; 1.29 |
| Cycle phase (fertile vs. post-fertile) | -0.01 | 0.52 | 0.03 | 0.97 | -1.04; 1.00 |
| Conception | -0.40 | -0.49 | 0.82 | 0.41 | -1.36; 0.55 |
| Parity | -0.05 | 0.43 | 0.13 | 0.90 | -0.89; 0.78 |
| Female rank | 0.28 | 0.21 | 1.33 | 0.18 | -0.13; 0.70 |
| Interaction: hindquarter luminance x |  |  |  |  |  |
| Cycle phase (fertile vs. prefertile) | 0.33 | 0.46 | 0.72 | 0.47 | -0.57; 1.22 |
| Cycle phase (fertile vs. post-fertile) | -0.65 | 0.40 | 1.61 | 0.11 | -1.44; 0.14 |
| *Conception* | *-0.96* | *0.52* | *1.82* | *0.068* | *-1.98; 0.07* |
| Parity | 0.00 | 0.37 | 0.00 | 1.00 | -0.73; 0.73 |
| Female rank | 0.07 | 0.20 | 0.33 | 0.74 | -0.32; 0.45 |
| Interaction: hindquarter redness x |  |  |  |  |  |
| Cycle phase (fertile vs. prefertile) | -0.53 | 0.51 | 1.04 | 0.30 | -1.53; 0.47 |
| Cycle phase (fertile vs. post-fertile) | -0.45 | 0.44 | 1.02 | 0.31 | -1.31; 0.41 |
| Conception | 0.56 | 0.64 | 0.88 | 0.38 | -0.70; 1.83 |
| Parity | 0.05 | 0.40 | 0.12 | 0.91 | -0.73; 0.83 |
| Female rank | 0.36 | 0.22 | 1.66 | 0.10 | -0.07; 0.78 |

**Figure 1.**
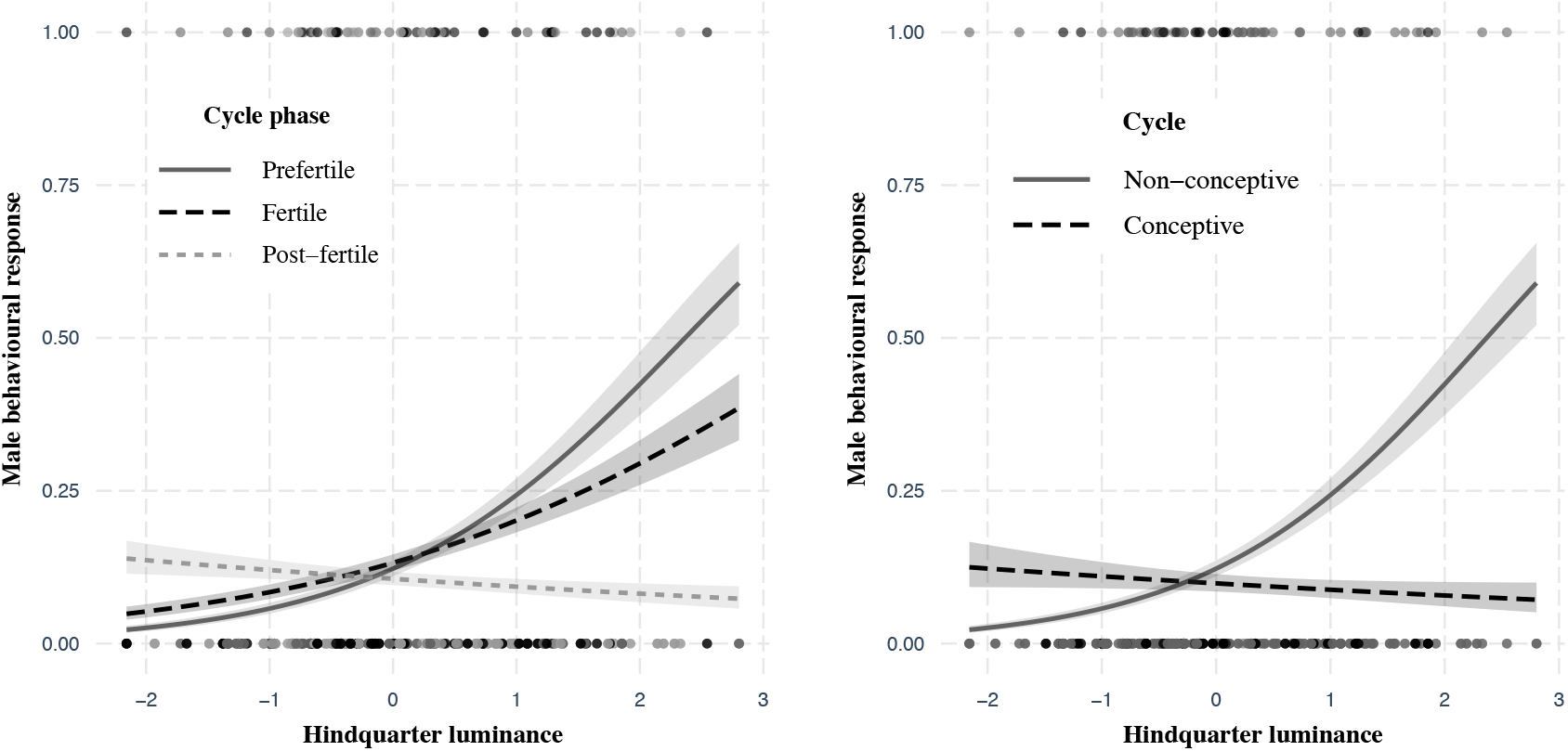
Interaction plot of the generalized linear mixed model exploring the relationship between male behavioural response, female hindquarter luminance (log) across cycle phases (A) and between conceptive vs. non-conceptive cycles (B).

Males were more likely to display sexual behaviours toward females that have solicited or interacted with them (ß = 2.99 ± 0.42 SE, P < 0.001, 95% CI = 2.15; 3.82, Table 3). Males were also more likely to display sexual behaviours toward females of lower parity (ß = -1.17 ± 0.53 SE, P = 0.026, 95% CI = -2.20; -0.14, Table 3) and of higher social rank (ß = 0.60 ± 0.28 SE, P = 0.031, 95% CI = 0.05, 0.15, Table 3). Male social rank was positively related to the occurrence of male sexual behaviours (alpha vs. beta: ß = -0.97 ± 0.37 SE, P = 0.010, 95% CI = -1.71; -0.24, alpha vs. gamma: ß = -1.45 ± 0.41 SE, P < 0.001, 95% CI = -2.26; -0.65, Table 3).

## Discussion

We examined male choice toward a non-sex-specific colourful ornament in a promiscuous primate species with traditional sex roles and demonstrated that males did not bias their sexual behaviours toward darker/redder females. Our results also showed that male Japanese macaques did not seem to use female colouration (luminance and redness of the face and hindquarters) to focus their mating effort on the most fertile period of the menstrual cycle, or during cycles that lead to conception, or on “good quality” mates.

The lack of effect of female colouration on male sexual behaviours -or in other words, the lack of male choice for darker/redder females-is surprising given that previous studies suggested a potential role of female colouration on male mate choice in Japanese macaques (face redness: Pflüger et al., 2014; face and hindquarter luminance and redness: Rigaill et al., 2019). While we investigated changes in male sexual behaviours toward group-member females varying in skin colouration and characteristics, Pflüger et al. (2014) investigated male responses toward two visual stimuli of an unknown female face. A red effect may have been over-estimated due to the binary choice, the lack of familiarity with the stimuli which significantly matters in communication (e.g., visual trait: Higham et al., 2011; odorant trait: Setchell et al., 2010; auditory trait: Townsend et al., 2011), or the lack of precise measure of attention such as tracking fine-scale eye movements (Hopper et al., 2020; Yorzinski et al., 2013). Our present study demonstrated that female colouration is unlikely to be a sexual signal since males did not appear to use this trait to discriminate when and with whom to mate to reproduce efficiently. Skin colouration may thus correlate with female characteristics without having a signalling function (correlation without causation). This highlights the importance of studying traits suspected to play a role in sexual communication and mating strategies from both signaller (what information may be contained) and perceiver (how the trait may affect behaviours) perspectives. Alternatively, we cannot completely rule out that mate choice for female colouration occurs in this species but that our data did not capture such effect. One may argue that males of this group may not rely – or need to rely – on this colouration to make their mating decisions as a result of the lower male-male competition due to the lack of peripheral males and the monopolization of mating from the alpha male. However, if female colouration indeed plays a role in male sexual strategies in this species where females vary in quality and female-female competition is high, one may have expected the alpha male to express mate choice as he has priority of access to mating. However, the alpha male did not appear to do so (see supplementary material).

In the absence of male mate choice for a female trait and of a clear ovulatory signalling in this species (Garcia et al., 2009; O’Neill et al., 2004; Rigaill et al., 2019), does this mean that males mate randomly? This is highly unlikely considering the energetic costs of extended sperm production, prolonged consortship, and male-male competition (Matsubara, 2003; Thomsen et al., 2006). While female colouration may not inform males about the probability of ovulation or female characteristics, this traits appears to be involved in the communication of pregnancy status (information content: Rigaill et al., 2015). Moreover, males have been found to reduce or avoid mating with pregnant females in populations facing higher energetic constrains (Fujita, 2010; Rigaill et al., 2015). However, whether males indeed use female colouration to avoid wasting energy in post-conceptive mating calls for further investigation. We also found in the present study that males were more likely to respond to female solicitations that could be used to apportion their mating effort toward receptive females vs. non-cycling females. Males may thus adapt their mating strategies according to female behaviour and to their access to mate by either intensifying mate guarding (higher-ranking males) or seizing a mating opportunity (lower-ranking males). Monitoring female behaviours would ultimately benefit to both higher- and lower-ranking males by balancing the costs and benefits of mate searching, direct competition, and mating. Alternatively, male sexual behaviours may be a by-product response of female mate choice which has been reported as an active reproductive strategy in Japanese macaques (Huffman, 1991, 1992; Soltis et al., 1997). We also found that males may bias their mate choice toward females at the beginning of their reproductive history and toward higher-ranking females. This may be related to parental investment as lower-ranking females and/or those toward the end of their reproductive history may have less energy to invest in future offspring or be less capable to balance the costs of reproduction with other costs (e.g., competition, health) than higher-ranking females and/or those at the beginning of their reproductive history (e.g., mandrills, Setchell & Wickings, 2006). More precise data on the interaction between female behaviours and characteristics and male mate choice is needed to better understand the process of male mate choice in Japanese macaques.

Studies of the role of non-sex-specific colourful ornaments displayed on feather or skin have been relatively limited in comparison with sex-specific colourful ornaments in both males and females. From this rather incomplete picture, it appears that male choice and preferences for female traits may evolve because of the costs associated with reproduction. One factor may be the cost of sexual signalling itself, when female ornamentations are costly to produce or maintain and thus impact the energy allocated to reproduction and ultimately, offspring quality and survival (Doutrelant et al., 2008, 2012; Griggio et al., 2009; Peters et al., 2011; Pilastro et al., 2003), although it is complicated to accurately assess such signalling costs. Another factor may be the shared costs of rearing offspring in socially monogamous species displaying extra-pair copulations (Griggio et al., 2005; Mahr et al., 2012; Torres & Velando, 2005). Under such conditions, both males and females would benefit from traits indicative of one’s fecundity to adjust their mating and parental effort according to the costs and prospective benefits of signalling to and of reproducing with a particular mate. Alternatively, male choice for a colourful trait has been also described in at least one promiscuous species with traditional sex roles. In the territorial agamid lizards, female colourful traits appear to convey information about receptivity: male choice may thus also evolve to maximize reproductive success when mating decisions are limited or constrained (LeBas & Marshall, 2000).

To our knowledge, our study is among the few to report a lack of male choice for female colouration (sockeye salmons: Foote et al., 2004; threespine sticklebacks: Wright et al., 2015). A possible explanation could involve the operational sex ratio. A female-biased operational sex ratio, i.e., when males are a limited resource, could favour female-female competition and ornamentation. However, in seasonal breeders, it is likely that the number of cycling, receptive, and defendable females within a group or location decreases rapidly during the mating season as a consequence of reproductive synchrony (Ims, 1990; Thompson, 2018). Male-male competition for the access to mates may thus strengthen which would select for male ornaments, especially in species where the cost of reproduction is heavier on females than males. Such scenario may explain the lack of male choice for female colouration in Japanese macaques. Females are indeed more likely to conceive at the beginning of the mating season and to invest high energy on gestation and maternal care while facing energetic stress from decreased food availability and thermoregulation (Fujita, 2010; Garcia et al., 2010, 2011). Thus, female colouration may only represent a correlated response of selection for ornamentation in males (Lande, 1980). The signalling function of male colouration and its role on mate choice in Japanese macaques is still unknown. However, evidence from other primate species suggests that male colouration can correlate with some aspects of male-male competition (geladas, *Theropithecus gelada*: Bergman et al., 2009; snub-nosed monkeys, *Rhinopithecus bieti*: Grueter et al., 2015; drills, *Mandrillus leucophaeus*: Marty et al., 2009; rhesus macaques: Petersdorf et al., 2017; mandrills: Setchell & Dixson, 2001) and may be used in female mate choice (rhesus macaques: Dubuc, Allen, et al., 2014; Dubuc et al., 2016; Waitt et al., 2003).

In conclusion, we found no clear evidence that female colouration (luminance and redness of the face and hindquarters) influence male mate choice in Japanese macaques. Female colouration does not appear to be a sexually selected trait conveying relevant information about intra-cycle, inter-cycle, and inter-individual differences in fecundity. However, future studies should investigate the potential role of both female and male colouration in the communication of reproductive status (e.g., cycling vs. non cycling, pregnancy), male-male competition, and female mate choice to draft any definite conclusions. Whether reproductive seasonality and reproductive costs influence the evolution of male mate choice and female traits remains an open question, especially in primates for which little data is available. Further studies focusing on the relationship between female/male colouration and male/female sexual behaviours and mate choice in species facing different reproductive constraint should help to improve our understanding of the role and evolution of non-sex-specific colourful ornaments in primates.

## Acknowledgments

The authors thank The Center for Human Evolution Modeling Research of the Primate Research Institute and M.A. Huffman for use of the primate facilities. We thank A. Chimènes for her help with the preparation of the digital photography. We also thank J. Duboscq and AJJ. MacIntosh for helpful discussion on the analyses and comments on an earlier version of this manuscript. This work was financially supported by the Centre National de la Recherche Scientifique (France) to CG.

