## supplementary material for "Does male mate choice select for female colouration in a promiscuous primate species?"

**Methods**

1) **Subjects**

| Female | Age (years) | Weight | Parity | Social rank |
|---|---|---|---|---|
| female 1 | 21.46 | 14.4 | 0.30 | 0.538 |
| female 2 | 14.48 | 10.0 | 14.48 | 0.231 |
| female 3 | 12.44 | 12.4 | 12.44 | 0.846 |
| female 4 | 10.22 | 10.5 | 10.22 | 0.154 |
| female 5 | 8.42 | 11.4 | 8.42 | 0.00 |
| female 6 | 7.37 | 9.0 | 7.37 | 1.00 |
| female 7 | 7.27 | 11.3 | 7.27 | 0.769 |
| female 8 | 6.55 | 9.4 | 6.55 | 0.615 |
| female 9 | 6.38 | 6.9 | 6.38 | 0.692 |
| female 10 | 5.54 | 10.0 | 5.54 | 0.308 |
| female 11 | 5.41 | 9.1 | 5.41 | 0.077 |
| female 12 | 5.37 | 8.8 | 5.37 | 0.462 |

**Table A**. Female characteristics.

2) **Variables**

| Name | Variable | Type | Levels or groups |
|---|---|---|---|
| Male behaviour | response | binary | 2 |
| Female behaviour | covariate | binary | 2 |
| Face luminance | predictor | continuous | - |
| Face redness | predictor | continuous | - |
| Hindquarter luminance | predictor | continuous | - |
| Hindquarter redness | predictor | continuous | - |
| Cycle phase | predictor | categorical | 3 |
| Conception | predictor | binary | 2 |
| Parity | predictor | continuous | - |
| Female rank | predictor | continuous | - |
| Male rank | covariate | categorical | 3 |
| Female identity | control | categorical | 12 |
| Male identity | control | categorical | 3 |
| Date | control | categorical | 73 |
| ***excluded from the models:*** | | | |
| Cycle number | predictor | continuous | - |
| Female age | predictor | continuous | - |

| | | | |
|---|---|---|---|
| Female weight | predictor | continuous | - |

**Table B**. Summary of the variables computed for the analyses

### 3) Models

glmer, with family = binomial(link="logit"), control = glmerControl(optimizer="bobyqa", optCtrl=list(maxfun=2e5))

**Null model** <- male behaviour ~ 1 + (1 | female identity) + (1 | male identity) + (1 | date)

**Face coloration model** <- male behaviour ~ (face luminance + face redness) * (cycle phase + conception + parity + female rank) + female behaviour +male rank + (1 | female identity) + (1 | male identity) + (1 | date)

**Hindquarter coloration model** <- male behaviour ~ (hindquarter luminance + hindquarter redness) * (cycle phase + conception + parity + female rank) + female behaviour +male rank + (1 | female identity) + (1 | male identity) + (1 | date)

**Face and hindquarter coloration model** <- male behaviour ~ (face luminance + face redness+ hindquarter luminance + hindquarter redness) * (cycle phase + conception + parity + female rank) + female behaviour +male rank + (1 | female identity) + (1 | male identity) + (1 | date)

### 4) Detailed results from each model

| | ß | SE | z-value | P | 95% CI |
|---|---|---|---|---|---|
| **Model face colouration** | | | | | |
| **Intercept** | **-1.19** | **0.54** | **-2.19** | **0.029** | **-2.25; -0.12** |
| Face luminance | 0.23 | 0.43 | 0.54 | 0.59 | -0.61; 1.07 |
| Face redness | 0.40 | 0.60 | 0.67 | 0.50 | -0.77; 1.57 |
| Cycle phase (fertile vs. prefertile) | -0.09 | 0.41 | -0.21 | 0.83 | -0.89; 0.72 |
| Cycle phase (fertile vs. post-fertile) | -0.37 | 0.42 | -0.88 | 0.38 | -1.19; 0.45 |
| Conception | -0.16 | 0.57 | -0.29 | 0.77 | -1.27; 0.95 |
| **Parity** | **-1.01** | **0.49** | **-2.08** | **0.037** | **-1.97; -0.06** |
| Female rank | 0.30 | 0.30 | 1.00 | 0.32 | -0.29; 0.90 |
| **Female behaviour** | **2.92** | **0.41** | **7.12** | **< 0.001** | **2.12; 3.72** |
| **Male rank (alpha vs. beta)** | **-0.97** | **0.37** | **-2.62** | **0.010** | **-1.69; -0.24** |
| **Male rank (alpha vs. gamma)** | **-1.44** | **0.40** | **-3.60** | **< 0.001** | **-2.23; -0.66** |
| Interaction: face luminance x | | | | | |
| Cycle phase (fertile vs. prefertile) | 0.44 | 0.46 | -0.97 | 0.33 | -1.33; 0.45 |
| Cycle phase (fertile vs. post-fertile) | -0.26 | 0.47 | -0.56 | 0.58 | -1.17; 0.65 |
| Conception | -0.51 | 0.46 | -1.12 | 0.26 | -1.41; 0.39 |
| Parity | 0.25 | 0.41 | 0.61 | 0.54 | -0.55; 1.04 |
| Female rank | 0.15 | 0.21 | 0.70 | 0.48 | -0.27; 0.56 |
| Interaction: face redness x | | | | | |

| | | | | | |
|---|---|---|---|---|---|
| Cycle phase (fertile vs. prefertile) | 0.21 | 0.53 | 0.38 | 0.70 | -0.84; 1.25 |
| Cycle phase (fertile vs. post-fertile) | -0.04 | 0.51 | -0.07 | 0.94 | -1.04; 0.96 |
| Conception | -0.41 | 0.48 | -0.85 | 0.40 | -1.36; 0.54 |
| Parity | -0.05 | 0.42 | -0.12 | 0.90 | -0.87; 0.77 |
| Female rank | 0.28 | 0.21 | 1.33 | 0.19 | -0.13; 0.69 |
| **Model hindquarter colouration** | | | | | |
| **Intercept** | **-1.20** | **0.53** | **-2.26** | **0.024** | **-2.24; -0.16** |
| Hindquarter luminance | 0.50 | 0.37 | 1.36 | 0.17 | -0.22; 1.23 |
| Hindquarter redness | -0.45 | 0.38 | -1.16 | 0.24 | -1.20; 0.31 |
| Cycle phase (fertile vs. prefertile) | -0.09 | 0.42 | -0.21 | 0.84 | -0.90; 0.73 |
| Cycle phase (fertile vs. post-fertile) | -0.25 | 0.40 | -0.63 | 0.53 | -1.03; 0.52 |
| Conception | -0.25 | 0.56 | -0.44 | 0.66 | -1.35; 0.86 |
| **Parity** | **-1.17** | **0.52** | **-2.23** | **0.026** | **-2.20; -0.14** |
| **Female rank** | **0.60** | **0.28** | **2.17** | **0.030** | **0.06; 1.15** |
| **Female behaviour** | **2.99** | **0.42** | **7.04** | **< 0.001** | **2.15; 3.82** |
| **Male rank (alpha vs. beta)** | **-0.97** | **0.37** | **-2.60** | **0.010** | **-1.71; -0.24** |
| **Male rank (alpha vs. gamma)** | **-1.45** | **0.41** | **-3.55** | **< 0.001** | **-2.26; -0.65** |
| Interaction: hindquarter luminance x | | | | | |
| Cycle phase (fertile vs. prefertile) | 0.33 | 0.45 | 0.72 | 0.47 | -0.56; 1.22 |
| Cycle phase (fertile vs. post-fertile) | -0.65 | 0.40 | -1.61 | 0.11 | -1.44; 0.14 |
| *Conception* | *-0.96* | *0.52* | *-1.83* | *0.068* | *-1.98; 0.07* |
| Parity | 0.00 | 0.37 | 0.00 | 1.00 | -0.73; 0.73 |
| Female rank | 0.07 | 0.20 | 0.33 | 0.74 | -0.32; 0.45 |
| Interaction: hindquarter redness x | | | | | |
| Cycle phase (fertile vs. prefertile) | -0.53 | 0.51 | -1.05 | 0.30 | -1.52;0.46 |
| Cycle phase (fertile vs. post-fertile) | -0.45 | 0.44 | -1.02 | 0.31 | -1.31; 0.41 |
| Conception | 0.56 | 0.64 | 0.88 | 0.38 | -0.70; 1.82 |
| Parity | 0.05 | 0.40 | 0.12 | 0.91 | -0.73; 0.83 |
| Female rank | 0.36 | 0.22 | 1.66 | 0.10 | -0.07; 0.78 |
| **Model face and hindquarter colouration** | | | | | |
| **Intercept** | **-1.14** | **0.58** | **-1.97** | **0.049** | **-2.28; -0.00** |
| Face luminance | 0.07 | 0.55 | -0.12 | 0.90 | -1.14; 1.00 |
| Face redness | 0.21 | 0.64 | 0.32 | 0.75 | -1.06; 1.47 |
| Hindquarter luminance | 0.53 | 0.45 | 1.16 | 0.24 | -0.36; 1.42 |
| Hindquarter redness | -0.47 | 0.43 | -1.10 | 0.27 | -1.32; 0.37 |
| Cycle phase (fertile vs. prefertile) | -0.08 | 0.46 | -0.18 | 0.86 | -0.99; 0.82 |
| Cycle phase (fertile vs. post-fertile) | -0.40 | 0.45 | -0.87 | 0.38 | -1.29; 0.49 |
| Conception | -0.10 | 0.62 | -0.15 | 0.88 | -1.31; 1.12 |
| **Parity** | **-1.34** | **0.64** | **-2.09** | **0.037** | **-2.59; -0.08** |
| Female rank | 0.51 | 0.31 | 1.65 | 0.10 | -0.10; 1.13 |
| **Female behaviour** | **3.11** | **0.45** | **6.89** | **< 0.001** | **2.22; 3.99** |
| **Male rank (alpha vs. beta)** | **-1.03** | **0.38** | **-2.68** | **0.007** | **-1.78; -0.28** |
| **Male rank (alpha vs. gamma)** | **-1.49** | **0.42** | **-3.57** | **< 0.001** | **-2.31; -0.67** |

| | | | | | |
|---|---|---|---|---|---|
| Interaction face luminance x | | | | | |
| Cycle phase (fertile vs. prefertile) | -0.22 | 0.54 | -0.40 | 0.69 | -1.28; 0.84 |
| Cycle phase (fertile vs. post-fertile) | -0.08 | 0.54 | -0.15 | 0.88 | -1.14; 0.98 |
| Conception | -0.34 | 0.54 | -0.63 | 0.53 | -1.39; 0.72 |
| Parity | 0.38 | 0.47 | 0.82 | 0.41 | -0.53; 1.29 |
| Female rank | 0.04 | 0.24 | 0.19 | 0.85 | -0.42. 0.51 |
| Interaction face redness x | | | | | |
| Cycle phase (fertile vs. prefertile) | 0.46 | 0.58 | 0.79 | 0.43 | -0.68; 1.61 |
| Cycle phase (fertile vs. post-fertile) | 0.25 | 0.56 | 0.44 | 0.66 | -0.85; 1.34 |
| Conception | -0.29 | 0.52 | -0.57 | 0.57 | -1.31; 0.72 |
| Parity | -0.09 | 0.48 | -0.19 | 0.85 | -1.03; 0.85 |
| Female rank | 0.35 | 0.23 | 1.51 | 0.13 | -0.10; 0.80 |
| Interaction hindquarter luminance x | | | | | |
| Cycle phase (fertile vs. prefertile) | 0.47 | 0.49 | 0.95 | 0.34 | -0.50; 1.44 |
| Cycle phase (fertile vs. post-fertile) | -0.51 | 0.46 | -1.11 | 0.27 | -1.40; 0.39 |
| *Conception* | *-1.18* | *0.60* | *-1.95* | *0.052* | *-2.36; 0.01* |
| Parity | -0.10 | 0.45 | -0.21 | 0.83 | -0.97; 0.78 |
| Female rank | 0.10 | 0.22 | 0.46 | 0.65 | -0.33; 0.54 |
| Interaction hindquarter redness x | | | | | |
| Cycle phase (fertile vs. fertile) | -0.47 | 0.57 | -0.83 | 0.41 | -1.58; 0.64 |
| Cycle phase (fertile vs. post-fertile) | -0.73 | 0.51 | -1.44 | 0.15 | -1.72; 0.27 |
| Conception | 0.48 | 0.76 | 0.64 | 0.52 | -1.00; 1.97 |
| Parity | 0.12 | 0.50 | 0.24 | 0.81 | -0.86; 1.11 |
| Female rank | 0.26 | 0.24 | 1.08 | 0.28 | -0.21; 0.72 |

**Table C.** Models parameters estimates (ß) ± adjusted standard errors (SE), z-value, probability (P), and confidence intervals (95% CIs) from each 4 candidate models. For each categorical comparison, the first level indicate baseline. Results for which CI does not include zero are presented in bold, trends are presented in italic.

### 5) Male rank and female colouration

glmer, with family = binomial(link="logit"), control = glmerControl(optimizer="bobyqa", optCtrl=list(maxfun=2e5))

**Face coloration model** <- male behaviour ~ (face luminance + face redness) * male rank + (1 | female identity) + (1 | male identity) + (1 | date)

**Hindquarter coloration model** <- male behaviour ~ (hindquarter luminance + hindquarter redness) * male rank + (1 | female identity) + (1 | male identity) + (1 | date)

**Face and hindquarter coloration model** <- male behaviour ~ (face luminance + face redness+ hindquarter luminance + hindquarter redness) * male rank + (1 | female identity) + (1 | male identity) + (1 | date)

| | ß | SE | z-value | P | 95% CI |
|---|---|---|---|---|---|
| **Intercept** | **-1.71** | **0.40** | **4.23** | **< 0.001** | **-2.51; -0.92** |
| Face luminance | -0.05 | 0.22 | 0.21 | 0.84 | -0.48; 0.39 |
| Face redness | 0.06 | 0.20 | 0.30 | 0.77 | -0.34; 0.46 |
| *Hindquarter luminance* | *0.37* | *0.21* | *1.81* | *0.071* | *-0.03; 0.78* |
| Hindquarter redness | -0.33 | 0.24 | 1.41 | 0.16 | -0.80; 0.13 |
| Male rank (alpha vs. beta) | -1.20 | 0.35 | 3.39 | 0.001 | -1.90; -0.51 |
| Male rank (alpha vs. gamma) | -1.22 | 0.34 | 3.57 | < 0.001 | -1.90; -0.55 |
| Face luminance x Male rank (alpha vs. beta) | -0.15 | 0.36 | 0.42 | 0.68 | -0.85; 0.55 |
| Face luminance x Male rank (alpha vs. gamma) | 0.07 | 0.36 | 0.20 | 0.84 | -0.63; 0.77 |
| Face redness x Male rank (alpha vs. beta) | 0.41 | 0.34 | 1.21 | 0.23 | -0.25; 1.06 |
| Face redness x Male rank (alpha vs. gamma) | 0.04 | 0.34 | 0.12 | 0.91 | -0.62; 0.70 |
| Hindquarter luminance x Male rank (alpha vs. beta) | -0.27 | 0.34 | 0.79 | 0.43 | -0.94; 0.40 |
| Hindquarter luminance x Male rank (alpha vs. gamma) | -0.32 | 0.33 | 0.94 | 0.35 | -0.97; 0.34 |
| Hindquarter redness x Male rank (alpha vs. beta) | -0.10 | 0.40 | 0.25 | 0.80 | -0.89; 0.69 |
| Hindquarter redness x Male rank (alpha vs. gamma) | 0.20 | 0.38 | 0.52 | 0.61 | -0.55; 0.95 |

**Table D.** Models parameters estimates (ß) ± adjusted standard errors (SE), z-value, probability (P), and confidence intervals (95% CIs). For each categorical comparison, the first level indicate baseline. Results for which CI does not include zero are presented in bold, trends are presented in italic.
